# A novel approach to develop wheat chromosome-specific KASP markers for detecting *Amblyopyrum muticum* segments in doubled haploid introgression lines

**DOI:** 10.1101/2021.09.29.462370

**Authors:** Surbhi Grewal, Benedict Coombes, Ryan Joynson, Anthony Hall, John Fellers, Cai-yun Yang, Duncan Scholefield, Stephen Ashling, Peter Isaac, Ian P. King, Julie King

**Affiliations:** Nottingham BBSRC Wheat Research Centre, School of Biosciences, University of Nottingham, Loughborough, UK; Earlham Institute, Norwich Research Park, Norwich, UK; Limagrain Europe, Clermont-Ferrand, France; USDA-ARS Hard Winter Wheat Genetics Research Unit, Manhattan, Kansas, USA; iDna Genetics Ltd., Norwich Research Park, Norwich, UK

**Keywords:** wheat, wild relative, *Amblyopyrum muticum*, SNP, KASP, introgression

## Abstract

Many wild relative species are being used in pre-breeding programmes to increase the genetic diversity of wheat. Genotyping tools such as single nucleotide polymorphism (SNP)-based arrays and molecular markers have been widely used to characterise wheat-wild relative introgression lines. However, due to the polyploid nature of the recipient wheat genome, it is difficult to develop SNP-based KASP markers that are codominant to track the introgressions from the wild species. Previous attempts to develop KASP markers have involved both exome- and PCR-amplicon-based sequencing of the wild species. But chromosome-specific KASPs assays have been hindered by homoeologous SNPs within the wheat genome. This study involved whole genome sequencing of the diploid wheat wild relative *Amblyopyrum muticum* and development of a SNP discovery pipeline that generated ∼38,000 SNPs in single-copy wheat genome sequences. New assays were designed to increase the density of *Am. muticum* polymorphic KASP markers. With a goal of one marker per 60 Mbp, 335 new KASP assays were validated as functional. Together with assays validated in previous studies, 498 well distributed chromosome-specific markers were used to recharacterize previously genotyped wheat-*Am. muticum* doubled haploid (DH) introgression lines. The chromosome specific nature of the KASP markers allowed clarification of which wheat chromosomes were involved with recombination events or substituted with *Am. muticum* chromosomes and the higher density of markers allowed detection of new small introgressions in these DH lines.

**Key Message:** A novel methodology to generate chromosome-specific SNPs between wheat and its wild relative *Amblyopyrum muticum* and their use in the development of KASP markers to genotype wheat-*Am. muticum* introgression lines.

## 1. INTRODUCTION

Bread wheat (*Triticum aestivum* L., 2n = 6x = 42, AABBDD) is one of the most widely grown and consumed crops worldwide. After two spontaneous interspecific hybridisation events (Dvořák et al. 1993; Marcussen et al. 2014; Pont et al. 2019), resulting in its allohexaploid genome, domestication and intensive breeding practices have reduced the genetic diversity available within and between modern bread wheat cultivars. The wild relatives of wheat, however, have a vast resource of untapped genetic variation that could be used to enrich and diversify the wheat genome. Recent studies have demonstrated the dramatic improvement in wheat-wild relative introgressions achieved through homoeologous recombination and genomics-based marker technologies (Qi et al. 2007; Tiwari et al. 2014; King et al. 2017; Grewal et al. 2018b; Cseh et al. 2019b; Xu et al. 2020).

Detection and characterisation of wild relative chromatin in a wheat background is an important requirement in wheat breeding programmes. Molecular markers provide a high-throughput and cost-effective way of achieving this and simple sequence repeats (SSRs) have been a popular marker system for the detection of wild relative introgressions because of their multi-allelic and co-dominant nature (Wu et al. 2006; Zhao et al. 2013; Fricano et al. 2014; Niu et al. 2018). However, with recent advances in Next Generation Sequencing (NGS) technologies and low-cost genome sequencing, single nucleotide polymorphism (SNP) markers are now the front-runner in the race to developing high-throughput genotyping platforms for marker-assisted selection (MAS) in crop breeding (Varshney et al. 2009; Rasheed et al. 2017). Exome-based sequencing of wheat varieties and wild relative species has resulted in a huge resource of SNPs (Winfield et al. 2012; Allen et al. 2013), which has been exploited to develop high-density SNP wheat genotyping arrays (Wang et al. 2014; Winfield et al. 2016; Allen et al. 2017). Wild relative introgressions have been detected in a wheat background using such wheat-based SNP arrays (Zhang et al. 2017; Zhou et al. 2018).

Genotyping of introgression lines is more efficient when using wild-relative genome-specific SNPs. The Axiom^®^ Wheat-Relative Genotyping SNP Array was developed (King et al. 2017) and used to detect introgressions from various wild species in a wheat background (Grewal et al. 2018a; Grewal et al. 2018b; King et al. 2018; Cseh et al. 2019a; Devi et al. 2019; Baker et al. 2020). Even though these genotyping arrays can be ultra-high-throughput and efficient, these SNPs cannot distinguish between homozygous and heterozygous individuals which limits their widespread use in crop breeding. Recently, whole genome and transcriptome sequencing have been used to develop genome-specific SNPs for *Lophopyrum elongatum* and tools such as high-resolution melting (HRM) markers and the Sequenom MassARRAY SNP genotyping platform, utilising these SNPs, were deployed for detecting *L. elongatum* introgressions in a wheat background (Lou et al. 2017; Xu et al. 2020).

The Kompetitive allele-specific PCR (KASP) platform has been demonstrated to be a flexible, efficient and cost-effective system for genotyping of introgression lines using wild relative genome-specific SNPs (Bansal et al. 2020; Grewal et al. 2020b; Han et al. 2020). However, hexaploid wheat’s polyploid genome makes it complicated to develop co-dominant interspecific SNPs. The first obstacle is the distinction between interspecific and an excess of homoeologous/paralogous SNPs found within the wheat genome. The second hurdle to overcome is the scoring of interspecific SNPs in segregating populations where the SNP has three homoeologous copies in the wheat genome. In such cases, it is difficult to differentiate between a heterozygous and a homozygous introgression line in a self-fertilized backcross population (Allen et al. 2011). Recently, Grewal et al. (2020a) addressed this problem by attempting to exploit interspecific SNPs with KASP assays that only had one copy of the template in the wheat genome. They reported wild relative genome-specific SNPs for ten species from the Amblyopyrum, Aegilops, Thinopyrum, Triticum and Secale genera using PCR-amplicon based sequencing of which, 620 were validated as chromosome-specific KASP markers in the wheat genome.

In this work, a more efficient bioinformatics-based approach was used to develop chromosome-specific KASP markers between *Amblyopyrum muticum* Eig. (2n = 2x = 14, TT) and bread wheat. Whole genome sequence of *Am. muticum* was generated using next generation sequencing and compared with the bread wheat cv. Chinese Spring, RefSeqv1.0 (Appels et al. 2018) high-quality reference genome sequence. Unlike previous work (Grewal et al. 2020a), only single-copy regions of the wheat genome were used for SNP discovery. A small subset of the SNPs were validated as KASP markers by genotyping a previously reported panel of doubled haploid wheat-*Am. muticum* introgression lines (King et al. 2019), increasing the density of KASP markers diagnostic for *Am. muticum*. The methodology reported here and the resulting KASP markers can be applied to other wheat-wild relative introgression studies and thus represents a valuable resource for the wheat research community.

## 2. MATERIALS AND METHODS

### 2.1 Plant Material

Four hexaploid wheat varieties (Chinese Spring, Paragon, Pavon76 and Highbury), three accessions of *Am. muticum* (2130004, 2130008 and 2130012; all obtained from the Germplasm Resource Unit at the John Innes Centre), three wheat-*Am. muticum* F_1_ lines (one with each accession of *Am. muticum*) and 67 doubled haploid (DH) wheat-*Am. muticum* introgression lines (King et al. 2019) were grown for leaf tissue collection and nucleic acid extraction.

All plants were grown in a glasshouse in 2L pots containing John Innes No. 2 soil and maintained at 18–25°C under 16 h light and 8 h dark conditions. Leaf tissues were harvested from 3-week-old plants, immediately frozen on liquid nitrogen and stored at -80°C until nucleic acid extraction.

### 2.2 Nucleic Acid Extraction

Leaf tissue (1.5 inch leaf segment cut into pieces) was harvested, frozen and lyophilised in a 2 ml 96 deep-well plate following which the samples were ground in the TissueLyser II (QIAGEN) using a steel ball in each well for 4-6 minutes at a frequency of 25 Hz. Genomic DNA for sequencing and genotyping was extracted according to the [Somers and Chao protocol](https://www.ars.usda.gov/ARSUserFiles/60701500/SmallGrainsGenotypingLaboratory/Protocols/wheat%20and%20barleyDNA%20extraction_original.pdf) (verified 10 September, 2021, original reference in Pallotta et al. 2003) from Step 2 onwards. For wild relatives with multiple accessions, the genomic DNA was pooled into one sample.

Genomic DNA extraction for generation of probes for genomic *in situ* hybridisation analysis, was carried using the above protocol with an additional step of purification with phenol/chloroform at the end.

### 2.3 Chromosome-specific SNP Discovery

DNA was isolated from *Am. muticum* accession 2130012, as described above, and a PCR-free library was prepared and sequenced on an Illumina HiSeq 2500 on rapid run mode to produce 101.86 Gb (∼16.50x coverage of *Am. muticum* assuming kew c-value genome size of 6.174 Gbp) of 250bp paired-end reads. To discover SNPs, the reads were mapped to the wheat reference genome assembly RefSeq v1.0 (Appels et al. 2018) using BWA MEM version 0.7.13 (Li 2013) with the -M flag. PCR duplicates were removed using Picard’s MarkDuplicates (DePristo et al. 2011) and alignments were filtered using SAMtools v1.4 (Li et al. 2009) to remove unmapped reads, supplementary alignments, improperly paired reads, and non-uniquely mapping reads (q<10). Variant calling was performed using bcftools (Li 2011) using the multiallelic model (-m). INDELs and heterozygous SNPs were removed and homozygous SNPs filtered using GATK VariantFiltration (DePristo et al. 2011). SNPs were retained if depth >= 5, allele frequency (AF) > 0.8 and quality score >= 30. To remove SNPs unsuitable for KASP assays, SNPs were removed if any other SNP was present within 50bp up or downstream. To prevent amplification of off-target regions in the genome, the SNP site along with 50bp up and downstream was aligned to RefSeq v1.0 using BLASTn (Camacho et al. 2009) and queries with any non-self-hits were discarded.

### 2.4 KASP Assay Design and Genotyping

To design a KASP™ assay, the flanking sequence of a SNP was fed through the PolyMarker application (Ramirez-Gonzalez et al. 2015) which aligned the query sequence to RefSeq v1.0 and provided two allele-specific primers and one common primer for each assay. A value of 1 in the ‘*total_contigs’* column of the output *Primers* file validated the query SNP to be in a single-copy region in the wheat genome RefSeq v1.0 assembly (Online Resource 2).

For genotyping purposes, three sets of KASP markers were used. Set 1 consisted of 150 chromosome-specific KASP markers previously reported to be polymorphic between wheat and *Am. muticum* (codes between WRC0001-1000; Grewal et al. 2020a). Set 2 consisted of 224 KASP assays designed to be tested on doubled haploid wheat-*Triticum urartu* introgression lines (codes between WRC1080-1308 and WRC1317-1393; Grewal et al. 2021). This is a subset of the 304 KASP markers developed in this study after 47 failed to amplify a PCR product and 33 were polymorphic between the parental wheat cultivars. Set 3 consisted of the new KASP assays designed in this study (codes between WRC1309-1316, WRC1394-1713, WRC1723-1872, WRC1894-1913, WRC1954-2113 and WRC2130-2169; Online Resource 2).

The genotyping procedure was as described by Grewal et al. (2020b). Briefly, the genotyping reactions were set up using the automated PIPETMAX^®^ 268 (Gilson, UK) and performed in a ProFlex PCR system (Applied Biosystems by Life Technology) in a final volume of 5 μl with 1 ng genomic DNA, 2.5 μl KASP reaction mix (ROX), 0.068 μl primer mix and 2.43 μl nuclease free water. PCR conditions were set as 15 min at 94°C; 10 touchdown cycles of 10 s at 94°C, 1 min at 65–57°C (dropping 0.8°C per cycle); and 35 cycles of 15 s at 94°C, 1 min at 57°C. Fluorescence detection of the reactions was performed using a QuantStudio 5 (Applied Biosystems) and the data analysed using the QuantStudio™ Design and Analysis Software V1.5.0 (Applied Biosystems).

### 2.5 Multi-colour Genomic *in situ* Hybridisation (mc-GISH)

Preparation of the root-tip metaphase chromosome spreads, the protocol for mcGISH and the image capture was as described in Grewal et al. (2020b). Briefly, genomic DNA from *T. urartu* (to detect the A-genome), *Aegilops speltoides* (to detect the B-genome), and *Aegilops tauschii* (to detect the D-genome) and *Am. muticum* were isolated as described above. The genomic DNA of (1) *T. urartu* was labelled by nick translation with ChromaTide™ Alexa Fluor™ 488-5-dUTP (Invitrogen; C11397; coloured green), (2) *Ae. speltoides* was labelled by nick translation with DEAC-dUTP (Jena Bioscience; NU-803-DEAC; coloured blueish purple), (3) *Ae. tauschii* was labelled with ChromaTide™ Alexa Fluor™ 594-5-dUTP (Invitrogen; C11400; coloured yellow) and 4) *Am. muticum* was labelled by nick translation with ChromaTide™ Alexa Fluor™ 546-14-dUTP (Invitrogen; C11401; coloured red). Slides were probed using 150 ng of *T. urartu*, 150 ng of *Ae. speltoides*, 300 ng of *Ae. tauschii* and 50 ng of *Am. muticum* labelled genomic DNAs, in the ratio 3:3:6:1 (green: blue: yellow: red). No blocking DNA was used. DAPI was used for counterstaining all slides. Metaphases were detected using a high-throughput, fully automated Zeiss Axio ImagerZ2 upright epifluorescence microscope (Carl Zeiss Ltd., Oberkochen, Germany). Image capture was performed using a MetaSystems Coolcube 1m CCD camera and image analysis was carried out using Metafer4 (automated metaphase image capture) and ISIS (image processing) software (Metasystems GmbH, Altlussheim, Germany).

## 3. Results

### 3.1 Generation of chromosome-specific SNPs

Alignment of *Am. muticum* WGS reads against the wheat reference genome RefSeq v1.0 and filtering for good quality uniquely mapped reads led to the identification of 38,137 SNPs in single-copy regions of the wheat genome (Online Resource 1). Fig. 1 shows the total number of SNPs found per wheat chromosome. In each homoeologous group the D genome chromosomes were found to have the most SNPs with *Am. muticum* with Chromosome 5D having the highest number of SNPs (3326) while chromosome 4A had the least (942).

**Fig. 1.**
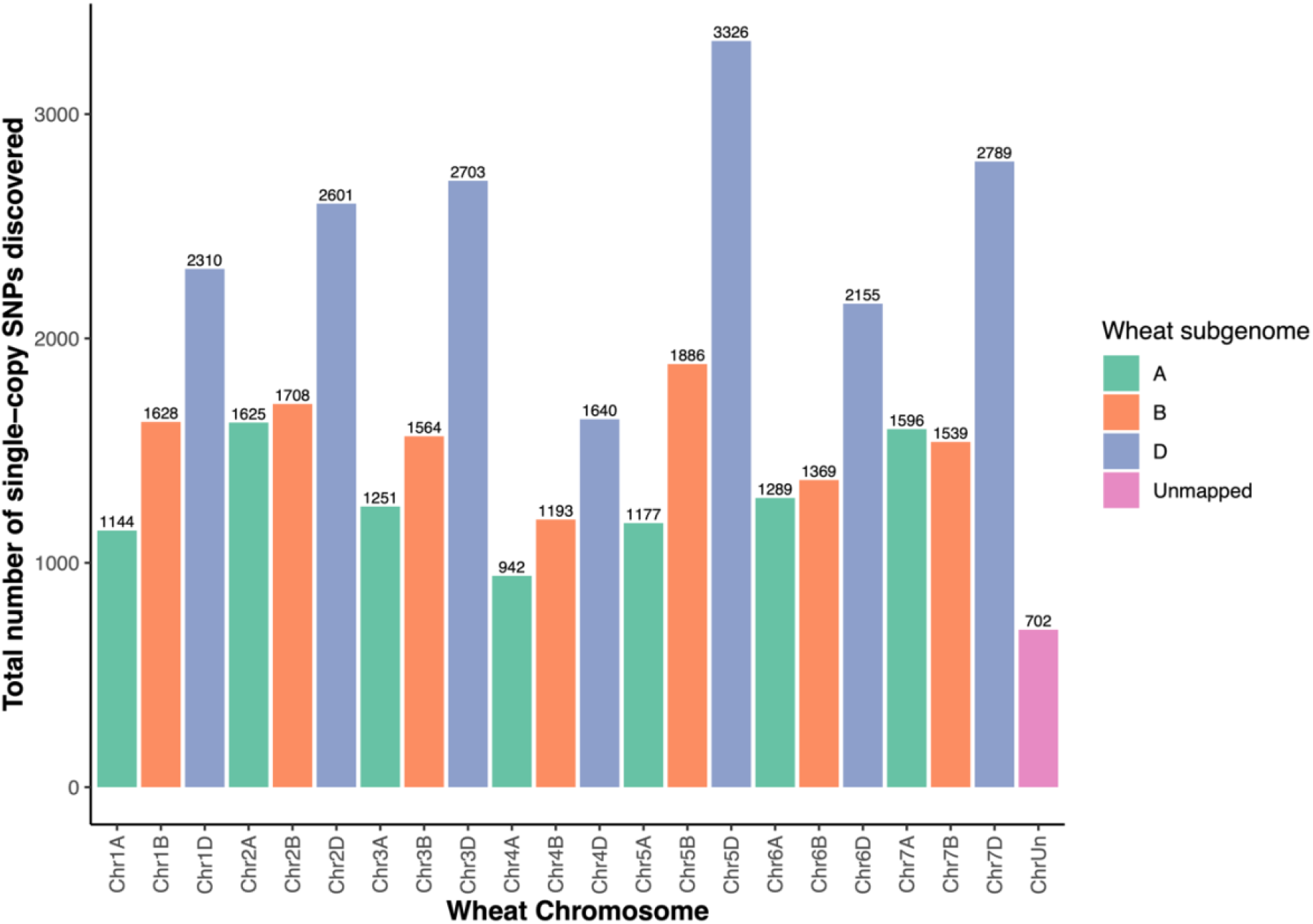
Plot showing the number of SNPs on each wheat chromosome, identified as polymorphic between hexaploid wheat and *Am. muticum* in single-copy regions of the wheat genome.

SNP density across 1,416 bins of 10 Mb each ranged from 0 (45 bins) to 221 SNPs (chr3D:600000000-610000000 Mbp). Fig. 2b depicts the range of SNP densities found across the wheat genome and bins with greater than 50 SNPs are shown in black. Of the 190 bins with more than 50 SNPs, 46 were present on A genome chromosomes, 19 on the B genome and 125 on the D genome. Most of the latter were found on the distal ends of D genome chromosomes (Fig. 2b).

**Fig. 2.**
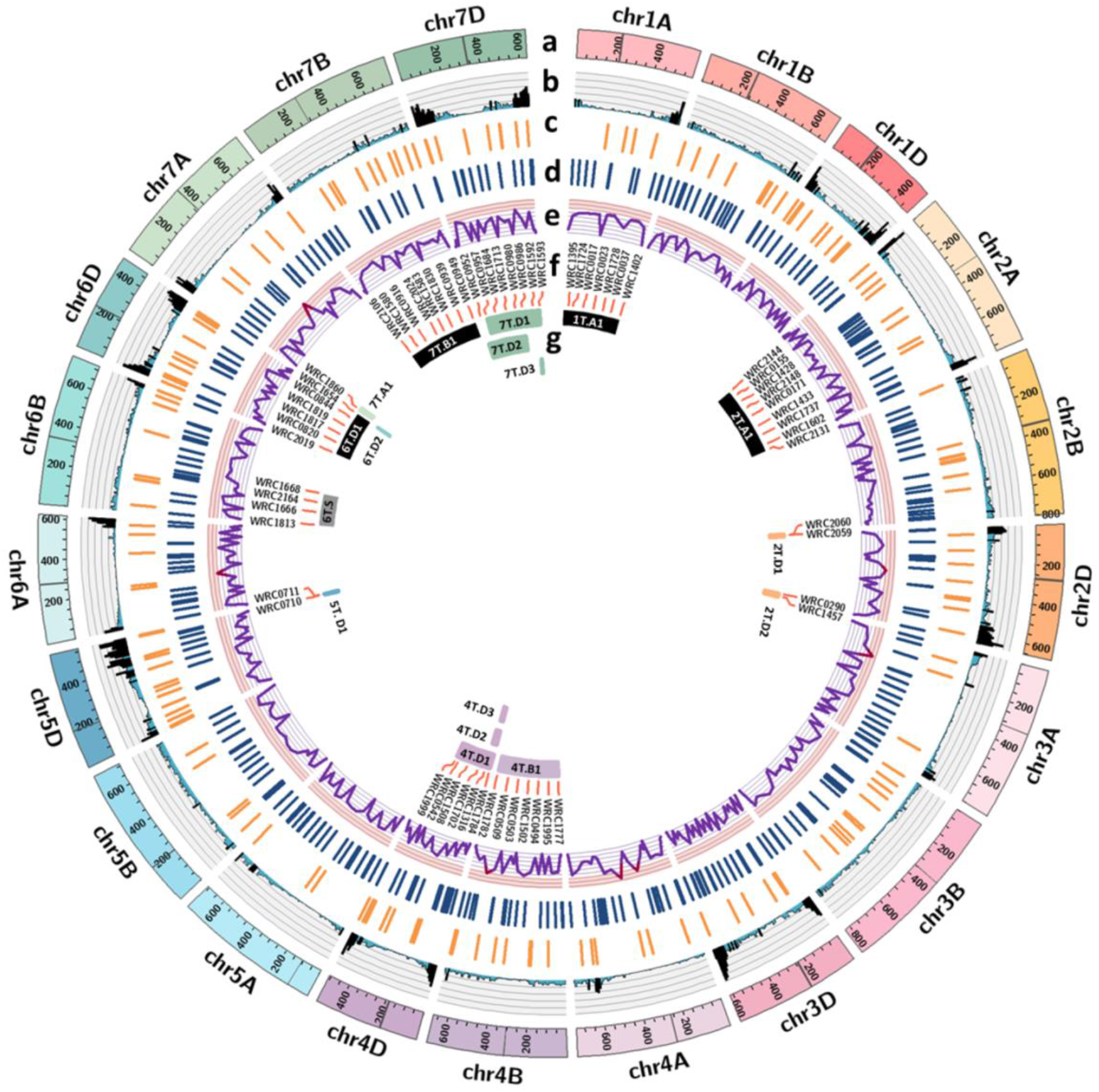
Circos plots of **a**. hexaploid wheat chromosomes (200 = 200 Mbp) with horizontal lines indicating position of centromere; **b**. SNP density in 10 Mbp bins (black = bins with >50 SNPs; starting at 0, each grid-line on the y-axis = 44.2 SNPs); **c**. position of functional chromosome-specific KASP markers in Set 1 and Set 2; **d**. position of functional chromosome-specific KASP markers in Set 3; **e**. distance between adjacent KASP markers on a wheat chromosome (red line = where distance between two markers >60 Mbp; starting at 0, each grid-line on the y-axis = 10.31 Mbp); **f**. a selection of KASP markers that detect all the introgression in the wheat-Am. muticum DH introgression lines; g. introgressions in the DH lines, coloured according to the corresponding recombinant wheat chromosome (black = disomic substitution, grey = disomic addition).

### 3.2 Chromosome-specific KASP marker development

Set 2 KASP markers previously developed and tested on *T. urartu* introgression lines (Grewal et al. 2021) were tested on wheat parental cultivars and *Am. muticum* accessions in this study. Of the 224 markers in this set, 194 (∼86.6%) failed to detect the *Am. muticum* allele, 17 (∼7.6%) were monomorphic between wheat and *Am. muticum* and 13 (∼5.8%) were found to be polymorphic between wheat and *Am. muticum* accessions used in this study. The positions on the wheat chromosomes of these 13 markers together with the 150 from set 1, previously developed and validated to be polymorphic between wheat and *Am. muticum* (Grewal et al., 2020a), are shown in Fig. 2c.

With the aim of having a KASP marker every 60 Mbp on a wheat chromosome, 698 new KASP assays were designed (Online Resource 2) in gap regions and tested on wheat, *Am. muticum* and three wheat-*Am. muticum* F_1_ lines in this study. Of these, 251 (∼36%) failed at the PCR stage, 49 (∼7%) did not amplify the *Am. muticum* allele, 22 (∼3.2%) were polymorphic within the wheat cultivars used as controls and 10 (∼1.4%) were monomorphic between wheat and *Am. muticum*. Of the remaining 366 KASP markers that were polymorphic between wheat and *Am. muticum*, 31 failed to detect the *Am. muticum* allele in the heterozygous state. Thus, 335 KASP markers were found to be functional and robust and their positions on the wheat chromosomes are indicated in Fig. 2d.

In total, 498 well-distributed, chromosome-specific KASP markers (Online Resource 3), polymorphic between wheat and *Am. muticum*, were used for downstream genotyping of introgression lines. Fig. 2e shows a line plot of the physical distance between these markers in wheat where each gridline of the y axis represents 10 Mb physical distance on a chromosome. The distance between the markers ranged from just 3 bases to ∼82.5 Mb with an average distance of 26 Mb. The average distance between the tip of the short arm and the first marker on the arm was 2.9 Mb while that from the last marker to the end of the long arm was 2.3 Mb. There were only seven instances where the gap between two KASP markers exceeded the desired 60 Mb and these are shown with a red stroke in the line in Fig. 2e. All these gaps were due to poor availability of SNPs within the desired bin as shown by the corresponding SNP density plot (Fig. 2b).

### 3.3 Validation of KASP markers through genotyping of introgression lines

The set of 498 chromosome-specific KASP markers, containing markers developed in previous studies and in this work, were used to genotype 67 DH wheat-*Am. muticum* introgression lines (King et al., 2019) along with parental wheat cultivars, *Am. muticum* accessions and F_1_ lines as controls. Previously, these DH lines were characterised using the Axiom^®^ 36K Wheat Relative Genotyping Array and multi-colour genomic *in situ* hybridisation (mcGISH) (King et al. 2017; King et al. 2019). The former technique provided information about what homoeologous group(s) from *Am. muticum* had introgressed into wheat and the latter identified the wheat subgenome(s) the *Am. muticum* segment(s) had recombined with and/or substituted, although, a drawback of the mcGISH technique is that it is unable to visually detect chromosome segments that are smaller than 18-20 Mbp.

In the current study, a homozygous introgression was detected through the presence of a homozygous *Am. muticum* allele called by KASP markers in the chromosome region the segment had recombined with or substituted (due to the absence of the wheat allele it had replaced) and through heterozygous calls by KASP markers that were present on homoeologous chromosomal regions in wheat (since these markers are also designed to be polymorphic with the introgressed *Am. muticum* segment but the corresponding wheat allele had not been replaced). Through genotyping of the DH lines with these chromosome-specific KASP markers we were able to validate most of the previous results and, in addition, identify the specific wheat chromosomes that the introgressions from *Am. muticum* had recombined with or substituted (Table 1). The markers helped in identifying specific cases of aneuploidy in some DH lines to support the GISH observations but also suggested disparities with previously reported results.

**Table 1.** Details of the type of introgression, its code (as indicated in Fig. 2g), and the wheat chromosome it had recombined with or substituted in each wheat-*Am. muticum* DH line as indicated through genotyping with chromosome-specific KASP markers. Observations about the chromosome constitution (deletions and aneuploidy) are also shown.

| DH Line name | Introgression Type: whole (W), arm (Telo), or recombinant (R) | Segment code<sup>a</sup> (A, B, D and T genomes represented as letters) | Wheat chromosome recombined with (if R) or substituted (if W) | Observations about chromosome constitution |
| --- | --- | --- | --- | --- |
| DH-1 | W | 6T.D1 | 6D |  |
| DH-6, 7, 8, 10, 11, 13, 339 | R | 2T.D2 | 2D |  |
| DH-15 | W, R | 2T.A1, 4T.B1 | 2A, 4B |  |
| DH-16 | W, R, R | 2T.A1, 4T.B1, 6T.D2 | 2A, 4B, 6D |  |
| DH-17, 18, 20, 21 | R, R | 4T.B1, 6T.D2 | 4B, 6D |  |
| DH-19 | R, R, W | 4T.B1, 6T.D2, 7T.B1 | 4B, 6D, 7B |  |
| DH-28 | Telo | 6T.S<sup>b</sup> | - |  |
| DH-29 | W | 7T.B1 | 7D |  |
| DH-62, 71, 74, 348 | R, R | 4T.D2, 7T.A1 | 4D, 7A |  |
| DH-63, 65, 66, 76, 77, 81, 341, 360 | R | 4T.D2 | 4D | 1D is missing in DH81, 7BL<sup>b</sup> missing in DH-360 |
| DH-83, 84, 85, 92 | R | 5T.D1 | 5D | 4BL is missing in DH-83 and DH-92 |
| DH-86, 91, 94 | R | 2T.D1 | 2D | 4 copies of 2D; interstitial deletion in 4A in DH-86 |
| DH-89 | R, R | 2T.D1, 5T.D1 | 2D, 5D | 4 copies of 2D; interstitial deletion in 4A |
| DH-96, 97 | R | 4T.D3 | 4D | 1DS is missing in DH-97 |
| DH-121 | R, R, R | 4T.D1, 5T.D1, 7T.D1 | 4D, 5D, 7D |  |
| DH-122 | R, R | 5T.D1, 7T.D1 | 5D, 7D |  |
| DH-123 | R | 7T.D1 | 7D |  |
| DH-124, 126, 128, 129, 131, 134, 137-139, 141, 144, 147, 355-357 | - | - | - | 1AL-1BL translocation<br>Tetraploid for 1A<br>Deletion of 1B not involved in 1A-1BL translocation |
| DH-161 | W | 1T.A1 | 1A | 7D is missing |
| DH-191, 192, 198, 202, 203 | R | 7T.D2 | 7D |  |
| DH-193, 195-197 | R, R | 7T.D2, 7T.D3 | 7D, 7D |  |

<sup>a</sup> From Fig. 2g
<sup>b</sup> S = short-arm, L = Long-arm

A subset of KASP markers that detect the *Am. muticum* introgressions present in these DH lines and the positions of these introgressions in the wheat genome are shown in Figs. 2f and 2g, respectively. In total, 17 different introgressions were found to be present in these lines, using this new set of chromosome-specific KASP markers, including 4 whole chromosome introgressions from 1T, 2T, 6T and 7T, a telocentric introgression of the short arm of chromosome 6T and 12 large and small segments from chromosomes 2T, 4T, 5T, 6T and 7T that had recombined with various wheat chromosomes (Table 1 and Fig. 2g).

### 3.4 Deviations/Differences from previous characterisation of DH lines

As mentioned above, KASP markers that detect the introgression on the recombinant chromosome result in a homozygous call for the *Am. muticum* allele. In Figs. 3a-c, which show genotyping results of some of the DH lines, these homozygous *Am. muticum* calls are indicated in green. Markers on homoeologous wheat chromosomes that also detect the same introgression give heterozygous calls that are shown in red. The three wheat subgenomes are represented in shades of blue and indicate the presence of the wheat allele for KASP markers in those chromosomal regions. Fig 3b and 3c also show that heterozygous calls identifying introgressions from *Am. muticum* chromosomes were also obtained on non-homoeologous chromosomes in wheat due to chromosome rearrangements within wheat that were not present in *Am. muticum* such as the 4/5/7 translocation (Devos et al. 1995; Dvorak et al. 2018).

**Fig. 3.**
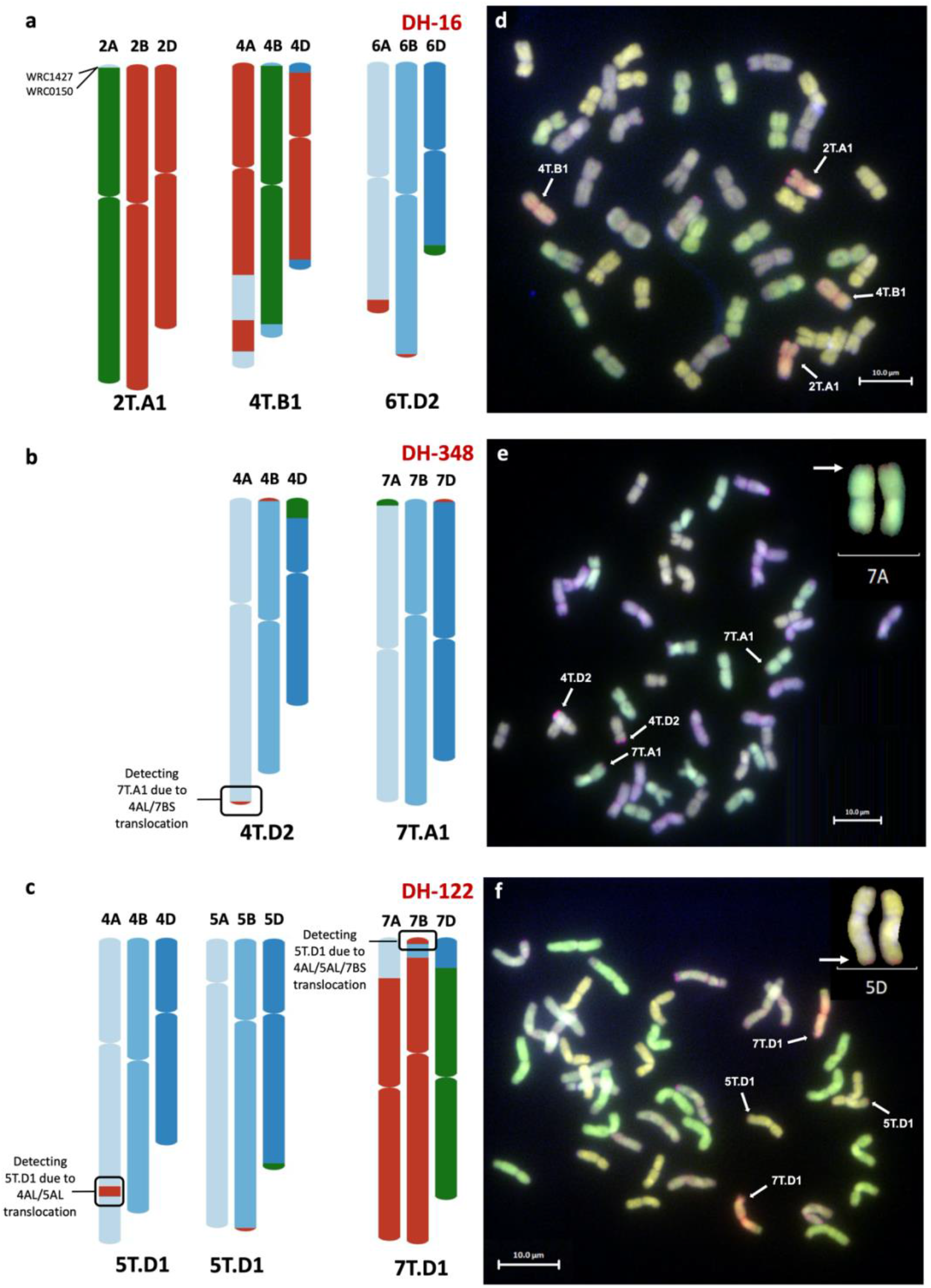
New small introgressions detected in wheat-*Am. muticum* DH introgression lines using chromosome-specific KASP markers. Graphical representation of KASP marker data detecting *Am. muticum* introgressions on wheat chromosomes in lines **a**. DH-16; **b**. DH-348 and **c**. DH-122. Shades of blue represent presence of homozygous wheat alleles, red indicates heterozygous calls and green indicates presence of homozygous *Am. muticum* alleles. McGISH analysis of root metaphase spreads validating marker data in **d**. DH-16; **e**. DH-348 and **f**. DH-122. Green indicates A-genome chromosomes, blueish grey indicates B genome, yellow indicates D genome and red indicates *Am. muticum* genome. White arrows point towards *Am. muticum* introgressed segments or chromosomes.

Previously, DH lines 15 and 16 were characterised as having two large introgressions from *Am. muticum* chromosomes 2T and 4T, both recombined with B genome chromosomes in wheat (King et al. 2019). Genotyping of these lines in this study showed that although chromosome 4T did recombine with chromosome 4B of wheat (4T.B1; Fig. 3a), chromosome 2T was introgressed as a whole chromosome that substituted a majority of chromosome 2A of wheat (2T.A1; Fig. 3a). KASP markers on the distal end of the short arm of chromosome 2A indicate that a very small segment of 2AS (∼12 Mbp) is potentially still present in these lines (Fig. 3a). However, GISH indicated that the 2T was potentially introgressed as a whole chromosome due to the presence of *Am. muticum* telomeric repeat signals on both ends of this introgression (Fig. 3d). If the 2AS segment had recombined with 2T or translocated onto another wheat chromosome, it would not be visible via GISH due to its small size.

The markers also showed that DH lines 16-21 had a small 6T segment (up to 10 Mb) on the distal end of 6DL (6T.D2; Fig. 3a) which was not previously detected by the Axiom array and is not visible by GISH. Genoytping analysis of four other DH lines 62, 71, 74 and 348, showed that in addition to the 4T.D2 segment, a very small segment (up to 20 Mb) from 7T was present at the distal end of 7AS (7T.A1; Fig. 3b) which had not been detected before in these lines. This very small segment on the distal end of chromosome 7AS was also detected by GISH in this study (Fig. 3e). The KASP markers were also able to detect another small segment (between 20-30 Mbp) from chromosome 5T in DH lines 121 and 122 (5T.D1; Fig. 3c). Due to its slightly bigger size, this *Am. muticum* segment can be viewed by GISH on the distal end of chromosome 5DL in DH-122 as shown in Fig. 3f.

Genotyping analysis of 15 DH lines (codes between DH 124-147 and DH 355-357), all shown previously to have a 1T introgression on chromosome 1A (King et al. 2019), showed that no introgression from *Am. muticum* was present in these lines. The absence of any call for the majority of the KASP markers on chromosome 1B indicated that these lines had lost the pair of 1B chromosomes but a small segment from the distal end of 1BL (∼50 Mbp) had been retained as indicated by the presence of wheat alleles for markers in this region. This new information potentially indicates that the translocation previously observed by mcGISH on a pair of 1A chromosomes and thought to be IT, was from chromosome 1BL.

## 4. Discussion

Previous studies have reported chromosome-specific KASP markers between wheat and *Am. muticum* (Grewal et al. 2020a) and other wild relative species (Grewal et al. 2020b; Grewal et al. 2021), which have been used for genotyping wheat-wild relative introgression lines. The objective of this work was to fill in the gaps with more KASP markers to increase the efficiency of genotyping by using an approach that involved faster SNP discovery and a more robust, chromosome-specific assay design than the ones reported in previous studies. In this work, we produced ∼38K SNPs between wheat and its wild relative *Am. muticum* in single-copy regions of the wheat genome and then converted some of these into wheat chromosome-specific KASP markers. In combination with previously designed chromosome KASP markers, a new set of well-distributed markers was obtained and used to re-genotype wheat-*Am. muticum* DH introgression lines (King et al. 2019) to validate the functionality of these markers as efficient genotyping tools and detect as many *Am. muticum* introgressions as possible.

A recently developed set of KASP markers (Set 2) was tested on *Am. muticum* accessions in this study but only 5.8% of the 224 assays were found to be polymorphic with wheat. This was as expected since this set of markers was originally developed to detect *T. urartu* introgressions in a wheat background (Grewal et al. 2021). When the 13 KASP markers were added to the 150 *Am. muticum* KASP markers developed during the original study (Grewal et al. 2020a), numerous gaps between markers were still present (Fig. 2c) preventing a uniform spread of markers able to detect *Am. muticum* introgressions across the whole of the wheat genome.

### 4.1 SNP Discovery

A major bottleneck at this stage was the lack of SNPs between wheat and *Am. muticum* that could be converted to KASP markers in regions that lacked an existing assay. With the advent of cheaper sequencing costs, it was possible to sequence the wild relative species to gain abundant SNPs for KASP assay design, some of which would be polymorphic between the species. However, in polyploid crops like bread wheat, it is challenging to generate chromosome-specific KASP assays able to distinguish heterozygous from homozygous individuals (co-dominant SNPs) and requires extensive validation (Allen et al. 2011; Allen et al. 2013; Grewal et al. 2020a; Makhoul et al. 2020). Thus, to avoid homoeologous SNPs which require a cumbersome KASP assay design process involving allele ‘anchoring’ for chromosome specificity (Grewal et al. 2020a), the approach taken here was to find SNPs in single-copy regions of the wheat genome using bioinformatic tools, thereby, resulting in ∼38K SNPs, each specific to a wheat chromosome (Fig. 1).

When the wheat genome assembly RefSeq1.0 was published (Appels et al. 2018), the authors reported the presence of 36,243 conserved subgenome orphan genes, which were defined as subgenome-specific genes found only in one wheat subgenome but having homologs in other plant genomes used in that study. They also reported the presence of 30,948 non-conserved orphan genes defined as either singletons or duplicated in the respective wheat subgenome, which did not have obvious homologs in the other subgenomes or the other plant genomes used in that study. Thus, the 38K SNPs in single-copy regions of the wheat genome generated in this work provided an excellent basis for marker generation although it was not known how many of these SNPs actually lay within orphan genes. In our previous work based on PCR-amplicon based sequencing and subsequent SNP discovery (Grewal et al. 2020a), only 18.2% of the 2374 SNP-containing sequences were found to be in single-copy regions of the wheat genome.

The D subgenome was found to have the most SNPs with *Am. muticum* (17,524), almost double those found with the A subgenome (9,024; Fig. 1). This could possibly have been because of more single-copy regions in the D subgenome than the A subgenome. However, the previous study suggested that the D subgenome had the least amount of orphan genes (19,523) compared to the A (22,496) and B (25,172) subgenomes (Appels et al. 2018). Another possibility could be that *Am. muticum* is more closely related to the progenitors of the D subgenome i.e., *Ae. tauschii* (McFadden and Sears 1946), which resulted in more *Am. muticum* sequence reads being mapped to the D subgenome chromosomes in turn producing more SNPs on the D subgenome. *Am. muticum* was previously classified under Aegilops species as *Aegilops mutica* Boiss. and like *Aegilops sharonensis* could be more closely related to D genome progenitors than B genome species (Marcussen et al. 2014). Early reports of homoeology of *Am. muticum* chromosomes suggested that T genome chromosomes pair with D genome chromosomes almost regularly in F_1_ from crosses of *Am. muticum* with D genome species (Jones and Majisu 1968) and recent reports have also shown that *Am. muticum* pairs more frequently with D and B genome chromosomes than with the A subgenome (King et al., 2017). Conversely, the increased number of SNPs with the D subgenome could be an indication of increased genetic diversity and sequence variation between the T and D genome species but not enough to prevent the sequences from being mapped onto the D subgenome.

SNP density analysis across 10 Mbp bins across the 21 chromosomes of wheat showed that SNP-dense regions (>50 SNPs per bin) were skewed to the distal ends of the chromosomes with a majority on the D genome chromosomes as shown in black in Fig. 2b. This is expected given that gene density on wheat chromosomes decreases towards centromeric regions (Appels et al. 2018; Brinton et al. 2020; Walkowiak et al. 2020; Przewieslik-Allen et al. 2021).

### 4.2 Development of KASP markers

A small portion of these SNPs (698) was selected to be to be converted into chromosome-specific KASP markers and added to the existing set of markers to provide a diagnostic marker for *Am. muticum* every 60 Mbp on a wheat chromosome. Of these, 48% (335) were validated as functional and robust, able to distinguish between heterozygous and homozygous introgression genotypes (Fig. 2d), while ∼36% (251) failed to amplify a PCR product. In the previous study involving development of chromosome-specific KASP markers (Grewal et al. 2020a) only 27% of the assays failed at the PCR stage. This could potentially be due to presence of sequencing errors or additional SNPs in the flanking sequence around the target SNP preventing efficient primer-binding in those sites and/or due to sub-optimal primer design. However, we reduced the percentage of assays that failed to detect the wild relative allele from 14.1% in the previous study to 7% in this study. The fact that *Am. muticum* is an outbreeding species and has an increased level of heterozygosity in its genome sequence could be contributing to the failure of KASP assays at the validation stage. If the target SNPs are present within the wild species and polymorphic for both the wild and wheat alleles, then it is possible that the DNA from plants used as controls for the *Am. muticum* accessions could be genotypes homozygous for the wheat allele or heterozygous at the target SNP.

We also observed 31 (∼4.4%) KASP assays that were polymorphic between wheat and *Am. muticum* but the heterozygous F_1_ reference genotypes wrongly clustered as homozygous with the wheat parents and thus, these KASP markers were deemed unsuitable for downstream genotyping of introgression lines. A previous study looking into this false SNP call assignments for some heterozygous genotypes suggested that artificial heterozygous DNA samples from parental lines, instead of natural heterozygote plants, can be used to identify false clustering (Makhoul et al. 2020).

This new set of KASP markers filled many of the gaps that existed between the markers developed using SNPs discovered through amplicon- or exome-based sequencing. The few regions where marker distances exceeded the desired 60 Mbp (Fig. 2e) were due to a lack of SNPs between wheat and *Am. muticum* in those chromosomal regions. Five out of these seven regions surrounded the centromeres of chromosomes 2D, 3A, 4A, 6A and 7A. As mentioned before, SNP density around centromeric and peri-centromeric regions is expected to be low (Brinton et al. 2020; Walkowiak et al. 2020) due to enrichment of sequence repeats and lower sequencing depths (Choulet et al. 2014; Wen et al. 2017).

### 4.3 Genotyping of *Am. muticum* DH lines

In total, 498 chromosome-specific KASP assays were used to genotype a set of 67 wheat-*Am. muticum* DH lines that had previously been characterised using a SNP array (King et al. 2019). Through KASP genotyping, these lines were shown to have 17 introgressions from the wild species including whole chromosomes and recombinant segments (Figs. 2f and g). Details of the parental lineage of each DH line has been published in the earlier study and as shown in Table 1, several sister DH lines inherited the same introgression segment from the wild species since they were progenies of the same BC_1_ plant.

KASP genotyping largely confirmed previous genotyping results of these DH lines but there were a few cases where the genotyping analysis provided new information which either negated previous results or highlighted new introgressions not observed previously. The latter included very small introgressions that were missed in the previous study due to a lack of markers in those regions and limitations of the GISH technique. After increasing the density of KASP markers available for identifying *Am. muticum* segments in this study and using the larger marker set for genotyping these DH lines, 3 new small introgressions were found: 6T.D2 on 6DL (up to 10 Mbp), 7T.A1 on 7AS (up to 20 Mbp) and 5T.D1 on 5DL (up to 30 Mbp; Table 1, Fig. 3a-c). The chromosome-specificity of the KASP markers allowed detection of the wheat chromosome that was involved in the recombinant chromosome or that had been substituted. Thus, it was observed that in DH lines 15 and 16, 2T.A1 was a whole chromosome that had replaced both 2A chromosomes rather than recombined with B genome chromosomes as previously reported. Where possible due to the size of the introgression, some of these results were validated by mcGISH in this work (Fig. 3d-f).

The chromosome-specificity of these KASP markers also allowed the detection of a number of wheat chromosome deletions in the DH lines as shown in Table 1. However, these were limited to the detection of homozygous deletions. These homozygous deletions included both whole wheat chromosomes or segments (both large and small) from the wheat chromosomes. In this context, one of the main observations involved the 15 sister DH lines (codes between DH-124 to 147 and DH-355 to 357) that showed that the pair of 1B chromosomes had been deleted in these lines expect for a small segment at the distal end of 1BL. We proposed that it was this 1BL segment that had translocated/recombined with a pair of A genome chromosomes, most likely chromosomes 1A. These lines also have 16 A genome chromosomes (King et al. 2019) and so it is possible that the pair of 1A-1BL recombinant chromosomes are present in addition to the pair of 1A chromosomes since the KASP markers at the distal end of 1A do not indicate the absence of any of the 1A wheat alleles.

## 5. Conclusion

Unlike previous work that relied on PCR-based amplicon sequencing (Grewal et al. 2020a), this method of generating SNPs between wheat and *Am. muticum* in single-copy regions of the wheat genome, made possible due to whole genome sequencing of the wild species, is rapid and allows for the development of chromosome-specific KASP assays. A variety of wild relative species are being used to increase the genetic diversity in hexaploid wheat. This approach can therefore be applied to other wheat wild relative species for SNP discovery, highlighting the need for greater investment in whole genome sequencing of these wild species. These KASP markers have greatly increased our capability to characterise, screen and identify both introgressions and wheat chromosomal aberrations in wheat-wild relative introgression lines. However, it is important to note that their efficiency is dependent on their density across the wheat genome and small introgressions existing between two KASP markers could have gone undetected. With the reducing cost of DNA sequencing, we envisage that the next improvement in characterisation of such introgressions, with the potential to give higher resolution, would be low-coverage whole genome resequencing of wheat-wild relative introgression lines.

